# Extreme high-diversity barcoded HIV library enables single cell multi-omics analysis of viral expression and fate determination

**DOI:** 10.1101/2025.01.14.632503

**Authors:** Tian-hao Zhang, Li Sheng, Xiangzhi Meng, Wenjin Lai, Yushen Du, Yuan Shi, Jing Wen, Masakazu Kamata, Sumit K. Chanda, Irvin SY. Chen, Ren Sun

## Abstract

HIV-1 persists as a global health challenge due to its ability to establish latent reservoirs that evade eradication by antiretroviral therapy (ART). This study introduces a high-diversity barcoded HIV-1 library combined with a novel peptide barcode system to enable single-cell multi-dimensional analysis of viral integration, transcription, splicing, and translation of distinct viral mRNA isoform. Our method revealed over 80 distinct HIV-1 splicing isoforms and uncovered significant temporal variation in mRNA diversity, driven by integration site and chromatin context. By linking HIV-1 5’UTR with peptide barcoding, we quantified the translation efficiency of alternative 5′ UTRs and demonstrated correlations with abundance of each viral mRNA isoform. Additionally, we showed that latency reversal agents, such as SAHA and Bryostatin, selectively reactivate proviruses depending on their genomic and chromatin context and leads to distinct viral RNA splicing program. This platform revealed the diversity of HIV-infected cells at levels of transcripts as well as at proteins, and subsequently the fate determination. The detailed understanding of the stochastic diversity and dynamic nature of HIV-1 latency, inform the development of targeted therapies to eliminate latent reservoirs.

## INTRODUCTION

Understanding the intricate dynamics of HIV-1 infection and the establishment of the HIV-1 reservoir is crucial for developing effective treatments and potential cures. Multi-omics methods provide a comprehensive approach to studying these complex processes. By capturing the full spectrum of molecular variations in individual cells that occur during HIV-1 infection, multi-omics analyses reveal the interplay between viral and host factors, identify biomarkers of disease progression, and uncover the mechanisms behind the establishment and maintenance of viral reservoirs. This holistic perspective is essential for facilitating the discovery of novel therapeutic targets and designing strategies to eradicate the latent HIV-1 reservoir, which remains a significant barrier to achieving a functional cure. Current single cell multi-omics have significantly contributed to the characterization of HIV-1 infection in vitro and in vivo. Various droplet-based or dilution-based methods can simultaneously characterize the viral transcriptome, flanking integration sequences, mutations in proviral sequences, along with phenotypes of the cells in which the provirus has been silenced^1–6^.

To date, these approaches have been used to investigate cell type and activity of the provirus, while neglecting the stochastics and dynamics of viral transcription and splicing. This is largely due to the scarcity of latently infected cells and the cost of single cell multi-omics methods. In this study, we describe a genetically barcoded HIV-1 library to profile viral integration, splicing and transcription at single cell resolution, but employing bulk-sequencing techniques. This eliminates the technical barriers to studying the behavior of single proviruses at a genomic scale and allows us to observe the viral splicing program with unprecedented resolution.

One arena where this methodology will be especially useful is the development of curative strategies for HIV-1 infection. Despite the continuous development of anti-retroviral therapies (ART), HIV-1 cannot be completely cured because it establishes latent reservoirs in patients. The latently infected host cells enter different degrees of dormancy wherein the proviruses are transcriptionally and translationally silent in the absence of an activating stimulus^7^. Various latency reversal agents (LRAs) have been developed to eliminate these reservoir^8–10^, including T cell stimulatory agents, kinase activators, and chromatin modifiers. However, in murine and non-human primate models of latency, as well as limited clinical trials, these regimens have largely proven to be ineffective due to incomplete penetrance. Additionally, there is great variability in their reactivation potential on an individual, cellular, and provirus basis. This heterogeneity in reactivation is thought to arise from complex interactions between chromatin structure and regulatory cis-acting elements near the site of proviral integration, the extracellular environment, as well as transcriptional regulators available in the cell. As a result, LRAs with different mechanisms will reactivate different subsets of proviruses, and the proviral integration sites are likely to play a major role in LRA efficay^11^. An additional issue is that LRA induced transcriptional bursting may not lead to virus production^12–13^. Thus, detailed profiling of the LRAs effect on viral transcription is needed.

Here we describe an innovative approach utilizing a genetically engineered, extreme high-diversity barcoded HIV-1 library to overcome existing technical barriers in single-cell multi-dimensional analysis. By linking viral integration, transcription, splicing and translation at single-cell level, we provided unprecedented insights into the stochastic and dynamic nature of viral life cycle. The methods enable the detailed profiling of proviral fate decisions in relation to integration sites and chromatin structure, as well as the biological consequences of alternative splicing. A viral 5’UTR library was constructed to illuminate the translational regulation on different viral mRNA isoforms.

Our findings reveal critical interactions between viral integration sites, splicing patterns and protein expression, significantly influencing the efficacy of latency reversal agents (LRAs) such as SAHA and Bryostatin. This work establishes a robust and cost-effective platform, reveals the functional diversity of HIV-infected cells at multiple levels, which expands our understanding of HIV-1 latency and reactivation, offering potential applications in developing curative strategies for HIV-1 infection.

## RESULTS

### Construction of an extreme high-diversity barcoded HIV-1 library

A genetically barcoded viral library was generated by insertion of 21-nucleotide random sequences between *env* and *nef* of HIV-1 NL-43 (Figure 1A). A 6-nucleotide translation insulator was placed downstream of the barcode to avoid interference with Nef translation. The barcodes include a cytidine every 3 nucleotides to avert arbitrary start codons. The average number of nucleotide differences between any two barcodes is 12 (Figure S1B) such that the possibility of 2 barcodes having less than a 3-nucleotide difference is 3x10^-6^. Nearly 2 million bacterial colonies were harvested and plasmid sequencing demonstrated that the library contains 2x10^6^ barcodes that are sequenced 3 or more times. The frequency distribution of the barcodes was uniform (Figure 1B) such that only 0.54% of them were found to be over-represented (>30 copies). We also determined barcode frequency after generating replication-competent virus and found it to be identical, within experimental error, to the frequency in the plasmid preparations from bacterial stocks (Figure 1C). This indicates the sequences of barcode do not impact viral replication.

**Figure 1.**
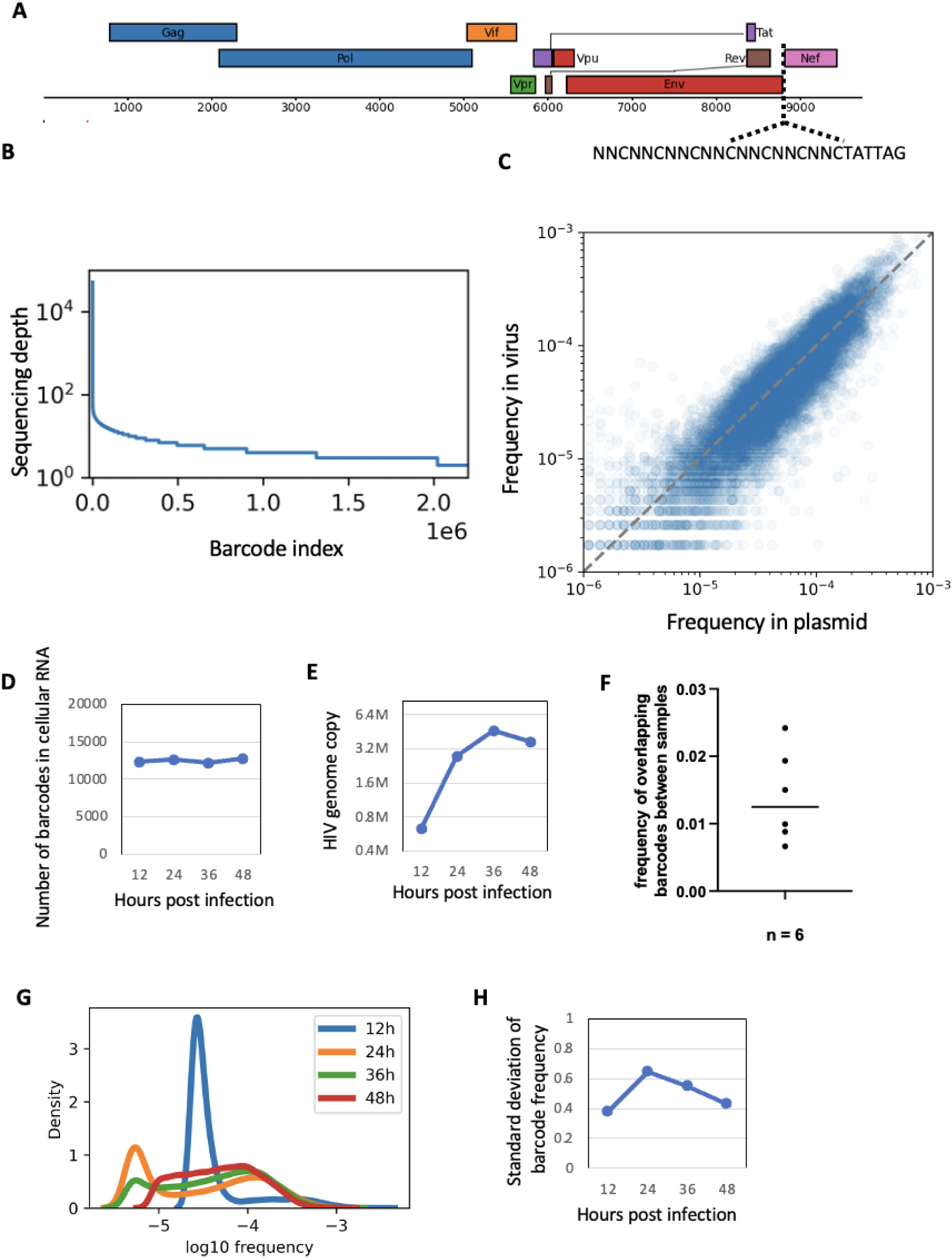
Characterization of the extreme high diversity barcoded HIV-1 library. **A)** The design of barcoded HIV. The barcode cassette was inserted between Env and Nef of a full-length HIV-1 genome. **B)** The frequency distribution of top 2 million barcodes in the plasmid library. The barcodes were sorted using their sequencing depth. Most barcodes were sequenced ∼ 10 times. **C)** The correlation between barcode frequency in the plasmid library and in the viral genomic RNA library produced in 293T cells. Each dot represents a barcode. **D-H)** Primary CD4+ T cells were infected with the barcoded viral library for 12h ∼ 48 hours at MOI 0.1. The cellular RNA was extracted, and the viral RNA barcodes were sequenced. **D)** The number of actively transcribing proviruses. The number of viral barcodes in the cellular RNA represents the number of viruses actively transcribing. **E)** The HIV-1 genome RNA copy in primary CD4+ T cells. The viral mRNA was quantified by RT-qPCR. **F)** The frequency of viral RNA barcodes overlaps with other samples. **G)** The histogram of provirus transcriptional activity. The x-axis is the relative frequency of viral RNA barcode in each sample, and the y axis is the occurrence of the barcode with certain frequency. **H)** The stand deviation of single provirus transcriptional activity in primary CD4+ T cells. The provirus transcriptional activity is quantified by calculating the relative frequency of each viral RNA barcode.

A simulation of an infection of 10^5^ cells showed that the probability of two cells being infected with a virus comprising the same barcode is less than 5% (Figure S1A). To validate this experimentally and to show that barcodes represent single infection events, 4 million CD3/CD28-activated CD4+ T cells were infected with barcoded HIV-1 at MOI 0.1. A protease inhibitor (Darunavir) was added to the cell culture to prevent multiple rounds of infection. The infected population was split into 4 equal samples and cellular viral RNA was harvested and sequenced at 12-hour intervals for 48 hours. We added a unique molecular identifier (UMI) to the viral specific reverse transcription primer, enabling absolute molecule quantification of each barcoded viral RNA. Less than 2.5% of barcodes were shared among the 4 samples, indicating that the majority represent independent infection events (Figure 1F). The number of barcodes detected in each sample indicated the number of proviruses actively transcribing viral RNA and remained constant in the 4 samples for the duration of the experiment (Figure 1D). However, the number of HIV-1 RNA copies measured by RT-qPCR increased during the first 36 hour of infection (Figure 1E). This indicates the increase of viral genomic RNA copies are not due to transcriptional activation of more proviruses, but the increased activity of already active proviruses. The observation also showed our method is sensitive enough to capture nearly all early transcriptional events, even when viral mRNA copy numbers per cell are low. Moreover, the frequency of the barcodes showed significant variation between sampling points (Figure 1G) indicating that provirus transcription was homogenous at 12 hours post-infection, and the variation is highest at the 24 hours post-infection (Figure 1H). The increase in transcription after the 12-hour time point was due to overexpression of a small portion of proviruses. In summary, the extremely high diversity of the barcoded HIV-1 library ensures that each cell will likely be infected by a virus comprising a unique barcode. This approach therefore enables quantifying barcodes to determine population diversity, and to use barcodes as a single cell labeling tool to study the behavior of individual proviruses.

### Barcoded virus enables single-cell profiling of viral splicing

To gain a more detailed understanding of the different transcriptional fates of HIV-1 proviruses, and to quantify the viral mRNA species in each infected cell, we developed a method to link barcode and splicing junction sequences from the same cells (Figure 2A). HIV-1 has 3 major classes of RNA products: unspliced (generating genomic RNA), single spliced (generating vif, vpr and vpu/env) and multi-spliced (generating tat, rev and nef). All splicing events take place upstream of the coding region, resulting in different 5’ UTRs (Figure 2B). Therefore, we only need the sequence of 5’UTR to infer the translation product of each mRNA isoform. To characterize splicing events, viral RNAs were first reverse-transcribed (RT) using a primer that adds a unique molecular identifier (UMI) immediately downstream of the barcode sequence. RT products were then amplified with PCR. The primers annealed to downstream of the UMI and upstream of the first donor site (D1). The UMI and 5’UTR were brought into proximity by self-circularization. Then, 3 set of linearization primers were used to generate short sequencing amplicons containing viral 5’UTR, UMI and the barcode. Each amplification product represented one of the 3 mRNA classes and was subjected to 150-bp paired-end sequencing. Thus, we captured more than 80 different splicing forms, including previously characterized, major forms (Figure 2B). Figure 2C illustrates the abundance of the different viral mRNA species in a representative cell. We recovered 28 different mRNA isoforms for all structural and accessory proteins in this one cell. Unspliced genomic RNA was the most abundant form, indicating this cell was producing virus. This method enables us to characterize the transcriptional and potentially translational status of each provirus.

**Figure 2.**
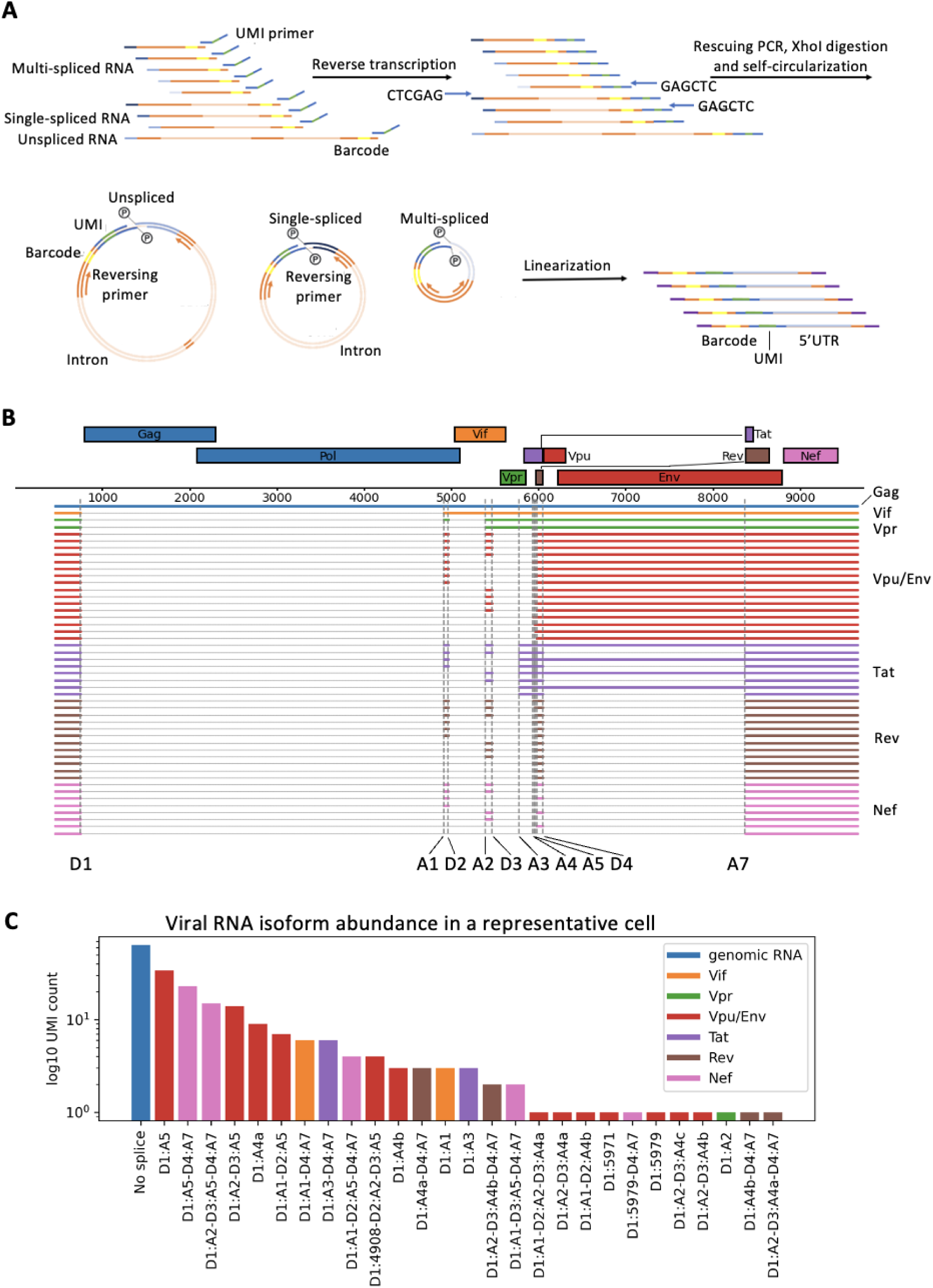
Sequences of single cell alternative splicing variants. **A)** Barcode - alternative splicing linkage workflow. Viral RNAs were reverse-transcribed (RT) using a primer that adds a unique molecular identifier (UMI) immediately downstream of the barcode sequence. RT products were then amplified with PCR. The primers annealed to downstream of the UMI and upstream of the first donor site (D1). A pair of XhoI cutting site was introduced on the PCR primers. The UMI and 5’UTR were brought into proximity by XhoI digestion and self-circularization. Then, 3 set of linearization primers were used to generate short sequencing amplicons containing viral 5’UTR, UMI and the barcode. **B)** The main forms of HIV-1 mRNA isoforms. The introns are shown as grey thin lines. The exons are solid color lines. Different viral ORFs share the same transcription start site but have different 5’UTR splicing and different translation start codon. **C)** Viral RNA isoform abundance in a representative cell. All viral RNA molecues in this panel have the same barcode but different UMI. The abundance of isoforms was quantified by counting the number of UMI.

### Provirus fate decision is a process of decreasing viral RNA heterogeneity

Next, we investigated the relationship between viral transcription and splicing by combining the splicing sequencing result with the viral RNA sequencing. We used the frequency of viral mRNA isoforms to calculate the entropy for each cell. This value represents the heterogeneity (diversity) of the viral mRNAs. The distribution of viral mRNA diversity decreased with the course of infection (Figure 3A). This could be explained by the bifurcation process during HIV-1 provirus fate decision. A provirus either goes into transcriptional burst, producing a large amount of viral RNA and entering the lytic cycle, or gradually shuts down its transcription and enters a latent state. In Figure 1, we showed that transcription of only a portion of viruses increased over time. In parallel, the mRNA diversity of individual viruses decreased over time, indicating that the provirus becomes more committed to transcribing a subset of viral mRNAs. To further characterize this phenomenon, we reconstructed the transcriptional dynamics of each viral open reading frame (ORF) in each cell (Figure 3B). Whereas the frequency of Tat, Vif, Vpr and Vpu/Env per cell increased over time, Rev/Nef decreased. This is consistent with previous reports of the 3-step transcription program^16^. The early viral genes, like Tat, Rev and Nef, are heavily spliced at the beginning of infection and decrease over time. Vif, Vpr and Vpu only need splicing of the D1-A4 intron and are expressed as the middle group. Late transcripts represent the full-length viral mRNA that can express structural proteins and that can be encapsidated. Despite the frequency change, we observed variations of the full-length RNA, Vif, Vpr, and Tat are decreasing over time, indicating a process of fate decision (Figure 3C). The frequencies of the RNA of these ORFs against total viral transcriptional activity were calculated. At the early time point, Tat, Vif and Vpr frequency correlated negatively with total viral RNA (Figure 3D). This orchestrates the function of Tat in promoting viral transcription. Rev is the only viral gene that is positively correlated with the within-cell diversity of viral mRNA, indicating an important role of RNA export in regulating mRNA diversity (Figure 3E).

**Figure 3.**
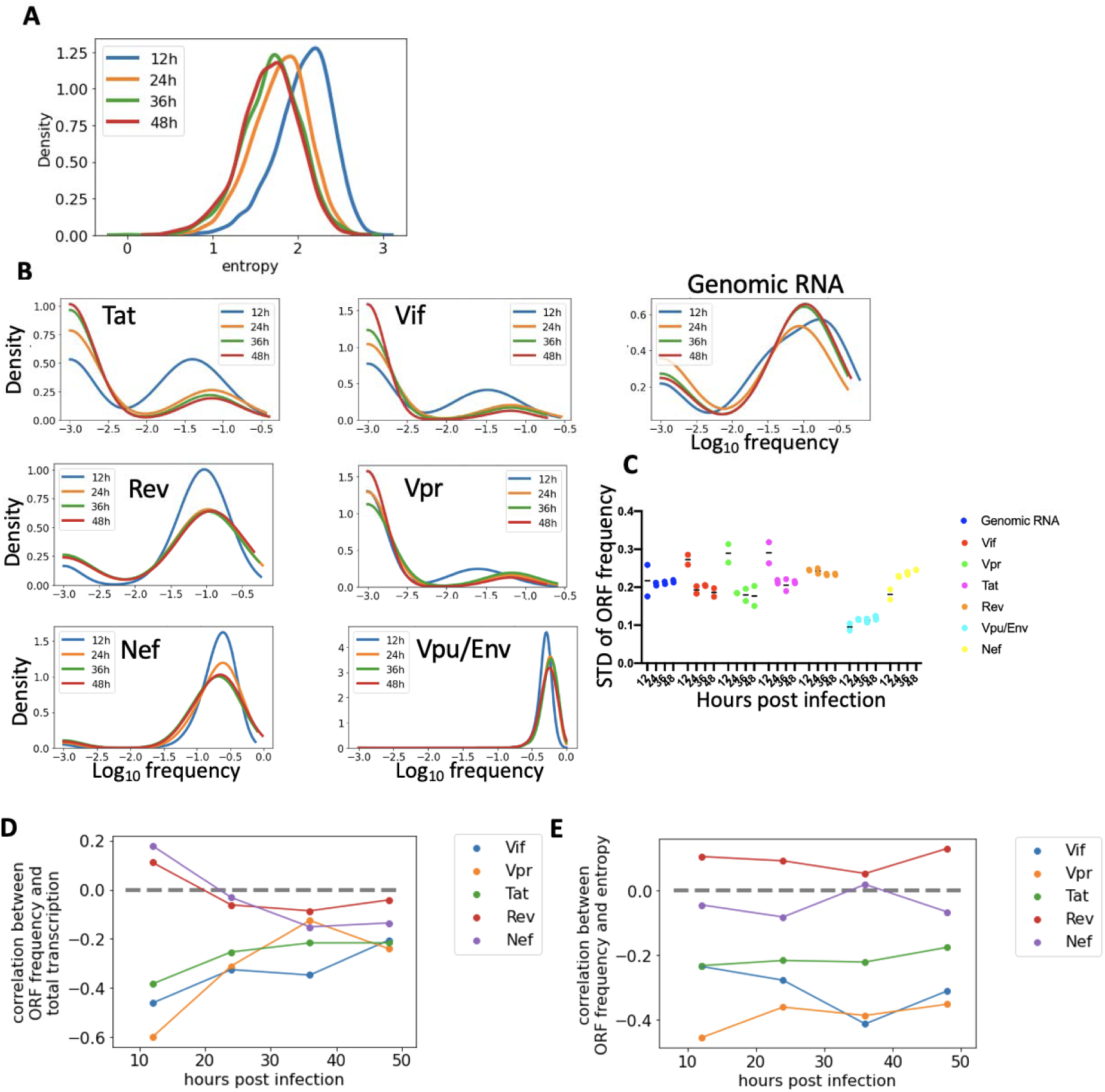
Single cell viral gene expression analysis. **A)** The histogram of the isoform frequencies’ entropy in each cell. Primary CD4+ T cells were infected with the barcoded viral library for 12h ∼ 48 hours at MOI 0.1. The cellular RNA was extracted, and the viral RNA isoform abundance was analyzed using the pipeline in Figure 2. The entropy of isoform frequencies represents the viral transcripts diversity. **B)** The histogram of viral gene frequency at different time point. The translation product of each viral RNA isoform was predicted according to its 5’UTR sequence. The relative frequency of each viral gene in each cell was calculated. **C)** The variation (standard deviation) of different viral ORF’s relative frequency at different time point. The data was calculated using the viral ORF frequency from panel B. **D)** The correlation between viral ORF’s relative frequency and total viral mRNA abundance in each cell. **E)** The correlation between viral ORF’s relative frequency and viral RNA’s entropy in each cell. The entropies are the same values calculated in panel A. Spearman’s correlation coefficients are shown in panel D & E.

The abundance and co-expression of different viral transcripts was investigated (Figure 4A). The abundance of genomic, Vif, Vpr and Tat mRNAs correlated positively, and these transcripts were usually co-expressed. Vpu/Env correlated negatively with the former group. The latter set of viral structural proteins, regulating the cellular environment for pyroptosis and virion budding, inhibiting the synthesis of cellular genes. Nef transcripts correlated negatively with all other genes and are indicative of tendency towards latency. Only a few viral mRNA molecules were heavily spliced into Nef mRNA. Rev transcripts were not correlated with any of the other transcripts (Figure 4B). Viral gene expression was summarized using principal component analysis (PCA) and plotted in 2 dimensions (Figure S2A). Proviral fate can be visualized by inferring a pseudotime based on gene expression using SlingShot (Figure S2B). The pseudotime correlated well with the real sampling time.

**Figure 4.**
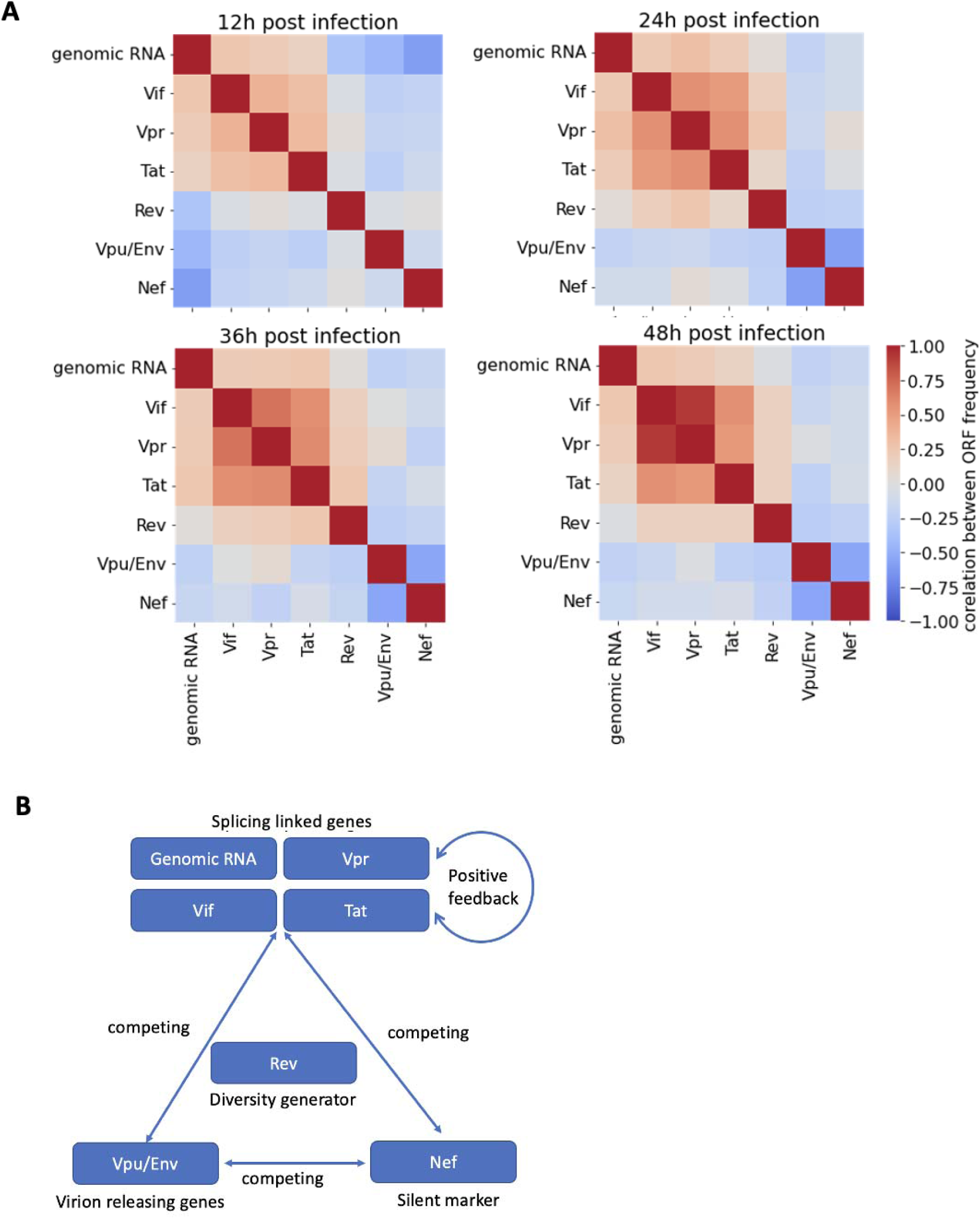
Correlation between viral ORF expression. **A)** The heatmaps showing Spearman’s correlation coefficients between different viral ORF’s relative frequency. The relative frequencies among genomic RNA, Vif, Vpr and Tat are positively correlated The relative frequency of Vpu/Env and Nef is negatively correlated with other genes. The relative frequency of Rev is not correlated with other ORFs. The correlations keep constant at different time points after the infection. **B)** The model of HIV-1 viral gene transcription regulation. Genomic RNA, Vpr, Vif and Tat are positively correlated. This splicing linked group competes with Vpu/Env and with Nef. These 3 groups of genes represent 3 different status of the viral life cycle, replication, releasing and latency.

### Barcode sequencing links viral integration sites with transcription

To study the effect of integration sites on viral transcription and splicing, we developed a barcode – integration site linkage sequencing method (Figure S3A). For each cell, we can link the proviral integration site with the viral barcode from paired end short read sequencing. We applied this method to HIV-1 infected primary CD4+ T cells and validated that the viral transcription mainly derived from integrated proviruses, but not the unintegrated or self-integration proviruses (Figure S3B/C/D).

The positions of the integrated proviruses were mapped onto a reference human genome. Consistent with previous publications, HIV-1 integration took place on all chromosomes (Figure S4A) and favored transcriptionally active region (Figure S4B)^14,15^. Moreover, integration sites were observed more frequently near active histone markers than repressive histone markers (Figure S5A). Proviruses were also more likely to be found in short interspersed nuclear elements (SINEs), and less likely to be in other endogenous retroviral LTR regions (Figure S5B). Proviruses that integrated in genic regions exhibited significantly higher transcriptional activity than those integrated in intergenic regions (Figure S4C). However, the transcriptional activity of the provirus and the activity of the host gene being integrated was negatively correlated (Figure S4D).

This may be due to promoter occlusion resulting from hyperactive host genes depleting nearby transcriptional factors. The transcriptional activity of proviruses also correlated with nearby histone modifications (Figure S4E). Actively transcribed proviruses were closer to activating histone markers than latent proviruses. The transcriptional activity of proviruses inside of repeat region SINEs were also higher than the proviruses outside SINEs. All these data confirm that local genomic structure and host transcriptional activity are important for viral transcription at a large scale and high resolution.

### The effectiveness of latency reversal agents depends on integration sites

To achieve a complete cure of HIV-1, latency reversal agents (LRAs) are being developed as part of “kick-and-kill” therapies that aim to activate the HIV-1 latent reservoir followed by destruction of virus-producing cells. Among the major categories of LRAs are histone deacetylase (HDAC) inhibitors and protein kinase C (PKC) agonist whose efficacy is likely to be affected by proviral integration sites. To evaluate this, resting CD4+ cells infected with barcoded HIV-1 were treated with the HDAC inhibitor Suberoylanilide Hydroxamic Acid (SAHA) or the PKC agonist, Bryostatin (a PKC agonist). CD3/CD28 beads were used as a positive control of T cell activation. Viral integration sites and splicing products were sequenced 24 hours after treatment. All treatments substantially increased the level of viral transcription (Figure 5A). However, when the transcriptional activity of each provirus was investigated by quantifying RNA barcode abundance (Figure 5B), we found that neither treatment reactivated all proviruses. Rather, only a portion of latent proviruses or already transcriptionally active proviruses were being reactivated. SAHA reactivated a greater proportion of proviruses than Bryostatin, while the average transcriptional activity of provirus activated by Bryostatin was more significant (Figure S6A, S6B). Both LRAs activated proviruses in genic and intergenic regions (Figure 5C), with SAHA being more effective in activating provirus in intergenic regions. Both drugs preferentially activated proviruses in proximity of activating histone markers, but SAHA also reactivated proviruses near the repressive marker H3K9me3 (Figure 5D). Lastly, we determined which viral mRNAs were being produced in response to the LRAs. Within-cell diversity of viral mRNA decreased after the LRAs treatment (Figure 6A) indicating that these compounds initiate fate decision programs and increase total transcriptional activity. PCA enabled us to visualize all proviruses according to their gene expression profiles. CD3/CD28-mediated activation significantly upregulated viral genomic RNA, and the phenotype of Byrostatin treatment was closer to CD3/CD28 than SAHA (Figure 6B). The viruses treated by SAHA and the combination drugs had higher amounts of Vif and Tat (Figure 6C), indicating larger phenotypic heterogeneity. The data suggest that optimization of LRA combination therapies will be most effective in developing an effective shock-and-kill strategy.

**Figure 5.**
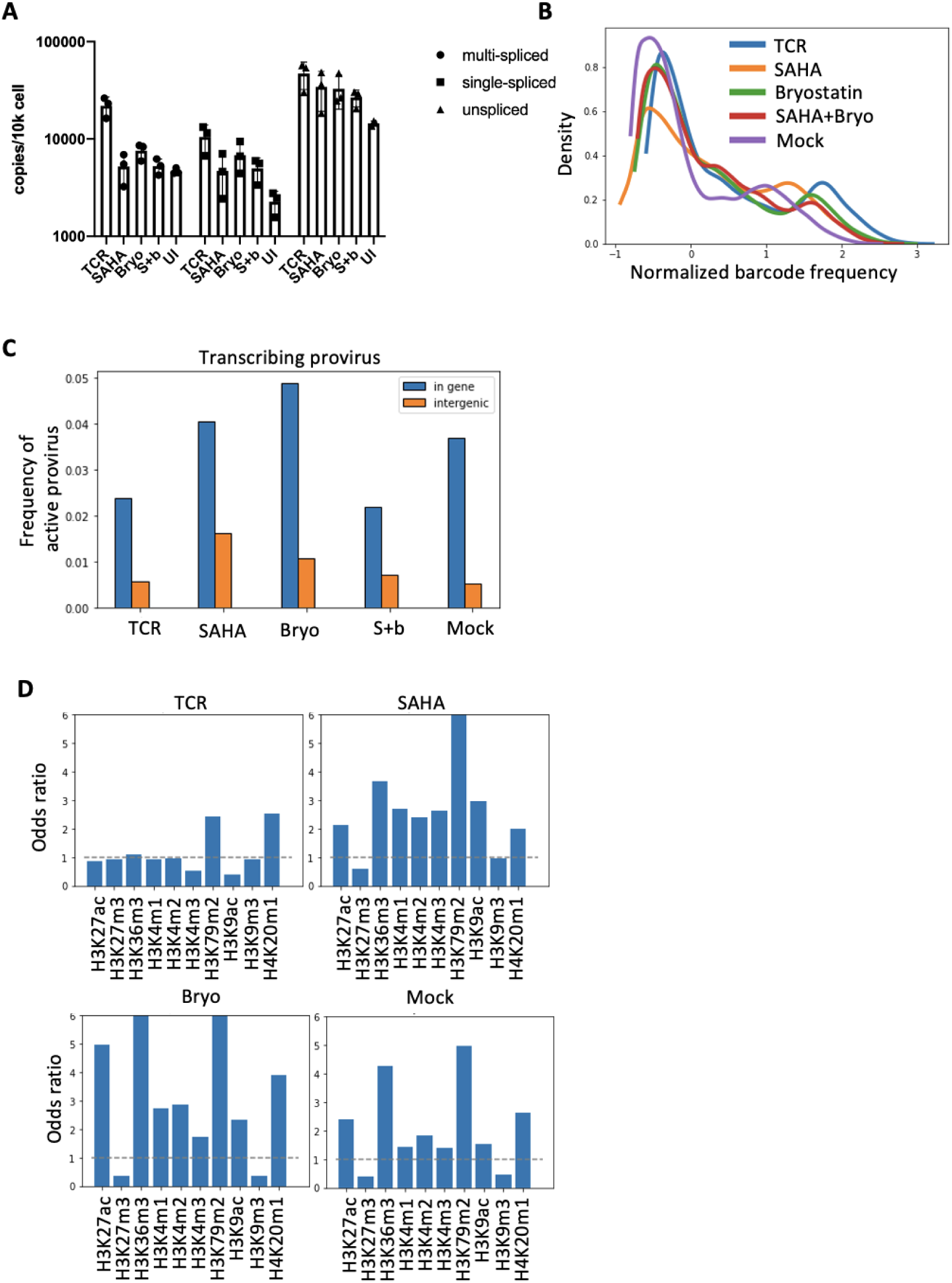
Effect of LRAs on different provirus. **A)** The copy number of viral mRNA after LRA treatment. The resting CD4+ T cells were infected with the barcoded virus library and reactivated with different LRAs. We used CD3/CD28 beads and recombinant IL2 to mimic TCR-induced activation. We also used SAHA and bryostatin (Bryo) to induce transcription activation. The combination group (S+b) used the same concentration of SAHA and bryostatin as the single drug treated groups. Untreated (UI) groups are used as control. 24 hours after reactivation, all cells were harvested for the analysis. Different groups of viral mRNAs were quantified using qPCR primers targeting different viral genes. Three replicates were performed. **B)** The histogram of viral RNA relative abundance in each cell. The x-axis represents the relative frequency of the viral RNA barcode in each cell. TCR-treated cells have the most high-frequency barcodes and the mock treated group has the least. **C)** The frequency of active provirus in different genomic regions. We used barcode integration site (IS) linkage sequencing to identify the integration site of the HIV-1 provirus in each cell and categorize the cells into “in gene” group and “intergenic” group according to the position of the integration site. The provirus is defined as active if its barcode is also observed in viral RNA sequencing. **D)** The odds ratio of actively transcribing provirus near certain histone markers. The number of integration sites within 5000bp of certain histone modification peaks were counted. Odds ratio larger than 1 represents a positively correlation between the integration site and the histone modification. ENCODE ChIP-seq dataset in CD4+ T cells were used.

**Figure 6.**
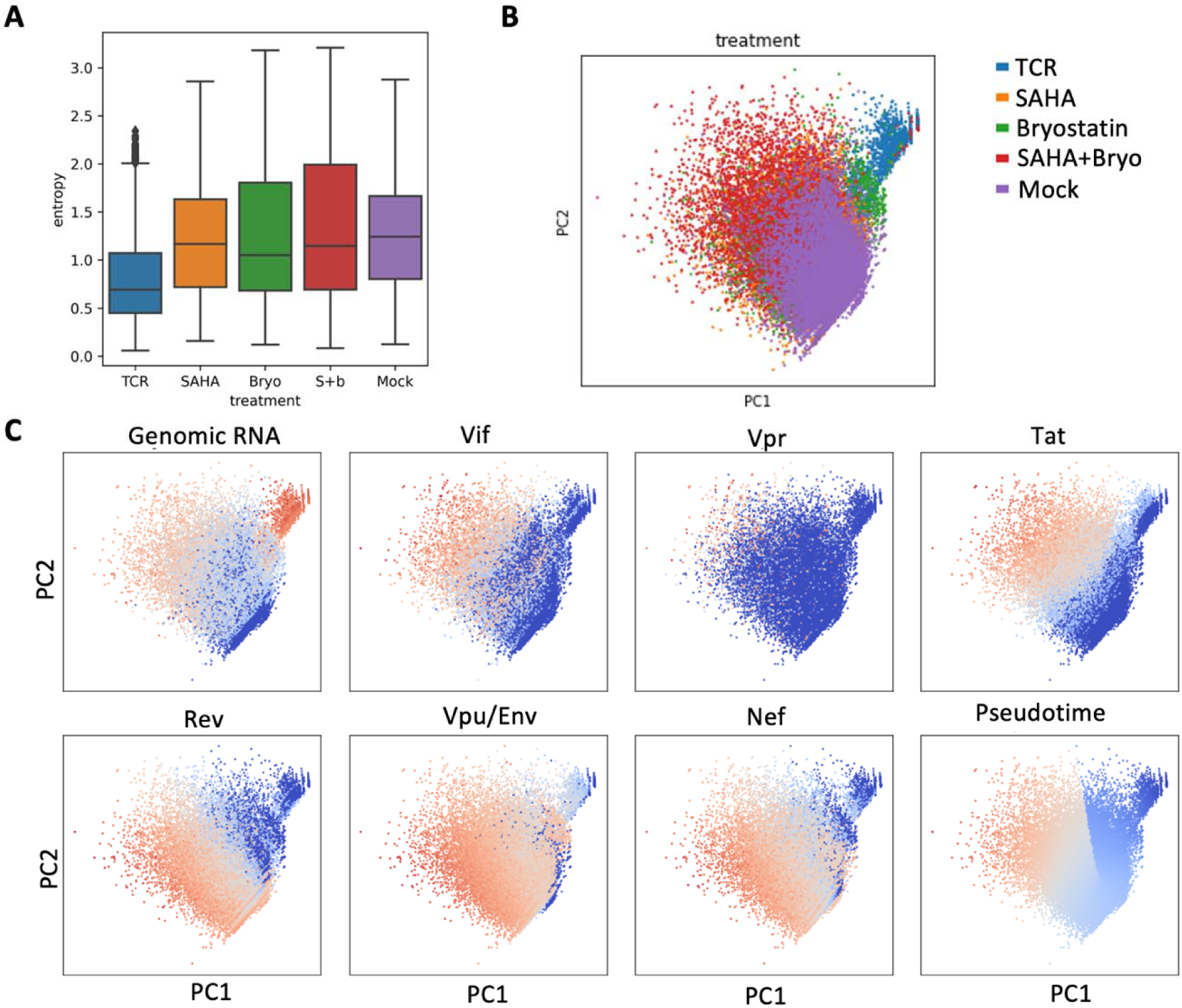
LRAs affect viral gene expression. **A)** The entropy of viral RNA frequencies in each cell with different LRA treated groups. The cellular RNA samples are from the experiment described in Figure 5. TCR-treatment reduced the viral RNA diversity. **B)** The PCA visualization of virus gene expression. The color code is identical to panel A. The relative frequencies of all isoforms were dimension reduced into 2 principal components. **C)** The PCA visualization of virus gene expression, colored by ORF’s abundance. Red color means higher gene abundance. Each panel is a different ORF. The last panel is the colored by pseudotime predicted by Slingshot. Red color means early pseudotime.

### The translation efficiency of HIV-1 5’UTR correlates with abundance of viral mRNA isoforms

HIV-1 genes share the same promoter and transcription start site. Their alternative splicing on 5’UTR resulted in different translation products and potential translation efficiency. To study the effect of different 5’UTR, we developed a peptide barcoding method (Figure 7A). Briefly, each HIV-1 UTR drives the expression of a GFP appended with a barcode of 18-19 amino acids . The peptide barcodes were designed to be effectively measured by mass spectroscopy to quantify the abundance of GFP translated from certain 5’UTR. We have included 66 HIV-1 5’UTR in the library.

**Figure 7.**
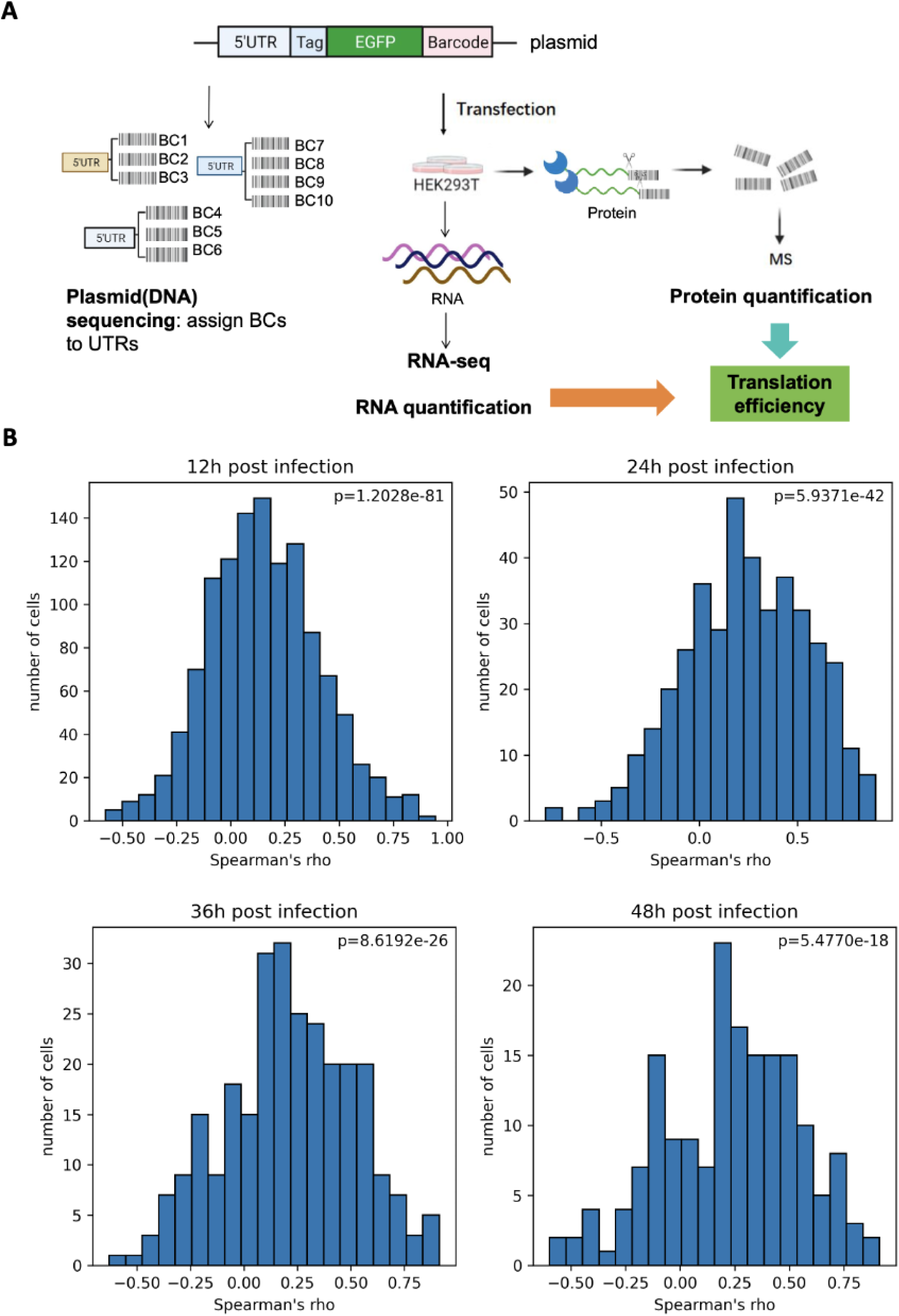
Quantification of the translation efficiency of different HIV-1 5’UTR using peptide barcode. **A)** The workflow of using peptide barcode to quantify the translation efficiency. An HIV-1 5’UTR library was linked with a barcode library fused with GFP. Each distinct 5’ UTR, indicated by different color, has ∼20 peptide barcodes. The library was transfected into 293T cells. The RNA was quantified by next generation sequencing and the peptide was quantified by mass spectroscopy. The translation efficiency of each 5’UTR was calculated by averaging the abundance of all linked peptide barcodes. **B)** The histogram of the Spearman’s correlation coefficients between the HIV-1 5’UTR translation efficiency and the abundance of its mRNA isoform in each cell. The HIV-1 infection was done as described in Figure 1. For each single cell (viral barcode), the correlation coefficient between log2TE (Translation Efficiency) and abundance of viral mRNA isoform was calculated.

Approximately 20 different peptide barcodes were linked to each 5’UTR. The translation efficiency was measured by averaging the relative frequency of all peptide barcodes linked to the 5’UTR and normalized to their frequencies in RNA.

We observed significant variation of the translation efficiency for different HIV-1 5’UTR. The translation efficiency among different viral transcripts’ strongest 5’UTRs are similar. But for each ORF, the translation efficiency of different UTR can vary by magnitudes. We also investigated the correlation between 5’UTR’s translation efficiency and its abundance of its mRNA (Figure 7B). We calculated the Spearman’s correlation coefficients for each mRNA isoform and found that they are significantly positively correlated. This data suggests the viral transcript isoforms with the highest abundance are the most efficient in translating viral proteins.

## DISCUSSION

Our study provides a significant advancement in understanding HIV-1 transcription and splicing by developing an extreme high-diversity barcoded HIV-1 library for single-cell multi-omics analysis. We provide a large dataset to further define that viral integration sites strongly influence transcriptional activity, with proviruses near active chromatin sites showing higher expression at a higher resolution. More importantly, we found that viral splicing patterns also vary based on integration sites, adding a new layer to the regulation of viral gene expression. Furthermore, different splicing products, viral mRNA isoforms, have different translational efficiency, which further increases the functional diversity among individual infected cells. Additionally, we showed that the response to latency reversal agents (LRAs) depends on the chromatin context, suggesting that different therapies may target different reservoirs. This methodology overcomes limitations of current single-cell technologies by effectively enrich viral RNA and capture the fine structure of viral splicing variants. It also offers unprecedented higher throughput at lower cost, allowing us to investigate how chromatin structure, transcription, and splicing jointly influence HIV-1 fate determination, including latency versus lytic infection. Our approach opens new questions, such as how to optimally target specific chromatin environments to fully reactivate latent reservoirs, providing critical information for developing more effective HIV-1 curative strategies.

The integration site of the virus can determine the accessibility of the host transcription and splicing machinery, thereby affecting the level and pattern of viral gene expression, and ultimately the outcome of the infection. We found the position of the HIV-1 integration site may affect the viral transcription and splicing, and consequently regulating the abundance of different viral proteins. This is achieved because the transcription and splicing machinery are not uniformly distributed in the host nucleus, and different intensity of splicing can lead to different viral gene expression patterns and functional variation, affecting virus fate decision. Studying the relationship between the integration site and alternative splicing can provide a more comprehensive understanding of the complex mechanisms of HIV-1 pathogenesis and may lead to the development of new treatment strategies that can specifically target and manipulate these processes. Testing drugs facilitating or inhibiting virus splicing together with LRAs may result in better coverage and efficiency of latency reversal.

SAHA remodels the histone modifications in the nucleus and may expose the beforehand dormant provirus. While Bryostatin induces the T cell activation signal and recruits many transcription factors into the nucleus. There are much crosstalk between two pathways. Eventually both lead to T cell activation and provirus transcription, but they have preference of different provirus according to the position of their integration sites.

Compared with current single cell multi-omics technologies, using an extreme high-diversity barcoded HIV-1 library allows analysis of ∼0.1 million cells for the cost of regular RNA sequencing. This is 50-fold higher throughput and 50-fold less costly than most single cell transcriptomic solutions. Our methodology therefore allows for greater throughput that could be useful in drug discovery campaigns or for use in animal models of HIV-1, SIV and SHIV infection.

The viral genome barcode method in this paper also has shortcomings. For example, it cannot characterize the phenotype of the infected cells. Additional cellular barcodes and microfluid system could be developed to circumvent this issue. Future studies can combine this method with droplet based single cell labeling or long read sequencing to generate a more comprehensive dataset for HIV-1 multi-omics.

The peptide barcode system was employed to quantify the translation efficiency of HIV-1 mRNA isoforms by linking specific 5′UTRs to unique peptide sequences that were designed to enable identification by mass spectrometry. Each peptide barcode served as a molecular signature, allowing precise measurement of protein translation products through mass spectrometry. By integrating this approach with high-diversity viral barcodes, we demonstrated that the abundance of viral mRNA isoforms correlates with their translation efficiency, revealing a hierarchy of translational output among different splicing variants. This result revealed another layer of regulation of viral expression, offering additional diversity of functions in each cell, fate determination of the infection, and subsequently related pathogenesis. This innovative use of peptide barcoding provides a powerful tool for dissecting the interplay between transcription, splicing, and protein production, offering critical insights into the regulation of HIV-1 gene expression and potential targets for therapeutic intervention.

## Methods

### Generation of barcoded HIV-1 library

The design of the barcoded virus is shown in Figure 1A. Two fragments were PCR amplified using the following primers: makeBC_F2(GGCTTGGAAAGGATTTTGCTATAANNCNNCNNCNNCNNCNNCNNCTATAAGATGGGTGGCAAGTGGTC) and makeBC_R2(GCTCCATGTTTTTCTAGGTC), makeBC_F1(CAGATCCATTCGATTAGTGAAC) and makeBC_R1(TTATAGCAAAATCCTTTCCAAGCC). The PCR generated 2 products of 343-bp and 176-bp respectively. The products were then purified using Invitrogen PureLink PCR purification kit and eluted in TE buffer (10mM Tris-HCl, 0.1mM EDTA, pH 8.0). The 2 fragments have a 24-bp overlapping region, so they were assembled using the NEB HiFi Assembly kit. The 495-bp assembled fragment was then amplified again using primers makeBC_F1 and makeBC_R2. The fragment was digested by restriction enzymes BamHI-HF and XhoI, and then purified. The pNL4-3 vector was also digested by these 2 enzymes and was purified by agarose gel electrophoresis. One ug of the insert fragment and 5ug of the vector was assembled using NEB HiFi Assembly kit. The assembled DNA was purified by ethanol precipitation with Pellet Paint NF co-precipitant. One ug of the purified DNA was transformed into the MegaX E. coli electrocompetent cells and plated to 20 15cm agar plates. The plates were cultured at 37°C overnight.

More than 0.5 million colonies were scratched from the surface of the plates. One mg plasmid DNA was extracted from the bacteria pellet. The barcode region on the plasmid was confirmed by Sanger sequencing. 20 ug of plasmid DNA was transfected into 20 million 293T cells using the Calcium phosphate transfection reagent. The virus library was harvested 48 hours after transfection. DNase I (40ng/mL) and MgSO4 (1mM) were added to the library to remove residual plasmid DNA from the supernatant. The barcoded virus library was aliquoted and frozen at -80°C for future use.

### Virus library infection

100 million HIV-1 free PBMC was obtained from anonymous donors. We use CD4 microbeads to isolate CD4+ cells from PBMC. The cells were then counted on a hemocytometer. The cells were cultured in RPMI-1640 with 10%FBS, 1% PenStrep, 10mM HEPES, 1mM sodium pyruvate, 0.1mg/mL Normocin, and 10U/mL recombinant h-IL2 with 5 million cells per mL density. 10uL CD3/CD28 beads were added to the culture for every million cells. 72 hours post activation, the CD3/CD28 beads were removed from the culture by a magnet and the cells were counted again. Virus worth 100ng p24 was mixed with 1 million CD4+ cells in the culture media and spinoculated for 90 minutes at 3000 rpm. After spinoculation, the cells were washed by fresh media and cultured with 10U/mL h-IL2 and 100nM Darunavir. The infected cells were harvested every 12 hours for subsequent assays.

### Sequencing library preparation

Total nucleic acid was purified using Qiagen DNA/RNA purification kit. The RNA was reverse transcribed using RT primer (CAAGTGCCTAGATCCTCGAGNNNNNNNNNNNNNNNNNNNNNCACTTGCCACCCAT

CTTATA) and purified by Invitrogen PCR clean-up kit. Here 21 consecutive random nucleotides serve as a unique molecular identifier (UMI) for each RNA molecule. For every infection, 3 types of the sequencing library were constructed. 1) Barcode amplicon library. 2) Barcode - integration site linkage library. 3) Barcode - splicing site linkage library.

For the amplicon library, the barcode region was amplified using primers CAAGTGCCTAGATCCTCGAG and GGCTTGGAAAGGATTTTGCTATAA. More than 10 million copies of viral cDNA were used as the template. This ensured sufficient coverage of most barcodes. The PCR product was confirmed using gel electrophoresis and purified by the PCR clean-up kit. I then used NEBNext Ultra II DNA library prep kit to make pair-end sequencing libraries. Around 50 million reads were sequenced for each sample.

The workflow of barcode integration site linkage sequencing was shown in Figure S3A. I used HinP1I (NEB) to digest the infected host genomic DNA. 63% of genomic fragments were at the length of 25 bp to 3000 bp, this ensures high circularization rate in the following steps. The digested DNA was then purified by PureLink PCR clean-up kit (Invitrogen). UltraII End-repair Module (NEB) prepared the DNA for ligation. A custom adaptor was annealed in the TE buffer (10mM Tris-HCl, 0.1mM EDTA, pH8.0). The sequence of the adapter’s reverse strand is TTGAGGTTTGCAGTTG. It has a 5-prime modification of a phosphorylation group, which facilitates TA ligation with the genome fragments. The 3 prime amino modification blocks the polymerase from adding nucleotides at its downstream, maintaining the L-shape conformation of the adapter.

Three consecutive phosphorothioate bonds at the 3-prime end stabilize the adapter, preventing it from enzymatic degradation. The forward strand of the adapter is ACCATCAACCCCGAATTCNNNNNNNNNNNNNNCAACTGCAAACCTCAAT. It anneals with the reverse strand and contains a 14-nucleotide UMI. 50pmol adapter was ligated to 1μg of fragmented genomic DNA. All ligated products were purified and amplified using 4 rounds of semi-nested PCR. All PCRs used the same reverse primer sequence: ACCATCAACCCCGAATTC. But the forward primer sequences anneal to different parts of the HIV-1 genome to increase the PCR specificity. They are in the order of F4 (AGTGAACGGATCCTTAGCACTTAT), F3 (CTCCTACAGTATTGGAGTCAGG), F2 (AGCCATAGCAGTAGCTGAGG) and F1 (GTACTCGAATTCGGGCTTGGAAAGGATTTTGCTATAA). All forward primers contain 3 consecutive phosphorothioate bonds at the 3-prime end, preventing the exonuclease activity of the polymerase, increasing the PCR specificity. Primer F3 and F2 are used with the reverse primer containing the 5-primer phosphorylation modification, to enable lambda exonuclease digestion after PCR, which can eliminate the product of unspecific amplification. The final PCR product was purified and digested by EcoRI-HF (NEB). This created two sticky ends on the DNA. 100 ng DNA was purified and subject to self-ligation in a 100μL reaction. The reaction used 2 units of T4 ligase (Invitrogen) at room temperature for 4 hours. The ligation efficiency was confirmed by quantitative PCR using primers ivF (ACACTCTTTCCCTACACGACGCTCTTCCGATCTTAGTCAGTGTGGAAAATCTCT), ivR (GAGTTCAGACGTGTGCTCTTCCGATCTTTTTGACCACTTGCCACCCAT) and synthetic standard templates. One thousand to 10 thousand copies of DNA per uL can be circularized. One third of the ligation product was used as the PCR template for the inverse PCR, using the same primers as the quantitative PCR. Phosphorothioate bond modification was used to increase PCR specificity. One tenth of the product was then subject to the final round of PCR, which adds Illumina sequencing adapters to the library.

The primers are AATGATACGGCGACCACCGAGATCTACACNNNNNNNNNNACACTCTTTCCCTACACGAC and CAAGCAGAAGACGGCATACGAGATNNNNNNNNNNGTGACTGGAGTTCAGACGTGTGC. N stands for indexing sequence to distinguish different samples. Around 10 million reads were sequenced for each library.

The workflow of the barcode - alternative splicing linkage sequencing was shown in Figure 2A. The near full length viral cDNA was amplified using primers CAAGTGCCTAGATCCTCGAG and ATCGATCTCGAGGCACGGCAAGAGGCGAGGG. Three consecutive phosphorothioate bonds at the end of the primers increased the PCR specificity. The amplified product was purified and digested by XhoI (NEB) for 2 hours. 1ng digested DNA was self-ligated in a 100uL ligation system. The reaction used 2 units of T4 ligase (Invitrogen) at room temperature for 4 hours. The ligation product was divided into 3 portions and subjected to 3 PCR. They all used the same forward primer (ACACTCTTTCCCTACACGACGCTCTTCCGATCTGGCTTGGAAAGGATTTTGCTATAA) which targets the upstream of the barcode region. But they use different reverse primers annealed to the downstream of the major splicing acceptor sites of the mRNA isoform families. For unspliced RNA, the reverse primer (GAGTTCAGACGTGTGCTCTTCCGATCTCTAGTCAAAATTTTTGGCGTACTCAC) anneals to the beginning of intron 1. For single spliced RNA, the reverse primer (GAGTTCAGACGTGTGCTCTTCCGATCTTCGTCGCTGTCTCCGCTTCT) anneals to the downstream of the A5 site. For multi-spliced RNA, the reverse primer (GAGTTCAGACGTGTGCTCTTCCGATCTCCCTCGGGATTGGGAGGTGG) anneals to the downstream of the A7 site. The PCR extension step only takes 20 seconds, so primers anneal to the downstream of the intended region would not have time for amplification and only 5UTR regions can be amplified. The cycle number was determined by the Ct number of a test run quantitative PCR, which only used 2uL of the ligation product. Finally, a 10-cycle PCR added Illumina sequencing adapters and indexes on the end of the library. Around 10 million reads were sequenced for each library.

All libraries were sequenced on the Illumina NovaSeq6000 platform using the PE150 setting.

### Peptide barcode library construction and translation efficiency profiling

The barcodes were designed with the following attributes: evenly distributed and easily detectable m/z range and hydrophobicity; a detectable length; and the inclusion of K in each barcode for TMT labeling, while avoiding M and R. The possible amino acids at each position of the barcode were manually assigned. All combinations were generated using Python. Out of a total of 248,832 combinations, hydrophobicity and m/z values were predicted by Prosit. Subsequently, barcodes with extremely similar m/z and hydrophobicity were filtered out, resulting in 96,000 barcodes for downstream experiments. As expected, these barcodes exhibited a well-distributed pattern in LC-MS/MS, facilitating identification and quantification in MS data analysis.

5’UTR, purification tag, vehicle protein, and DNA oligonucleotides encoding the corresponding barcodes were sequentially linked in the pLVX plasmid for transfection. To facilitate the detection of transfection efficiency, the vehicle protein was designed to be enhanced green fluorescent protein (EGFP). A protease cleavage site was inserted at the junction to enable the detachment of the barcodes and EGFP. At specific time points following transfection, cells were harvested for RNA and protein extraction. The RNA level encoding the barcodes was quantified using Illumina sequencing. The amino acid barcodes were digested from the EGFP with protease and quantified with LC-MS/MS. By comparing the quantities of RNA and amino acid barcodes, the regulatory effect of the 5’ UTR on translation can be assessed.

### Sequencing data analysis

The sequencing data were all analyzed by custom python codes and available upon request. Related sequencing data were deposited at under BioProject PRJNA970831.

---

**Figure S1.**
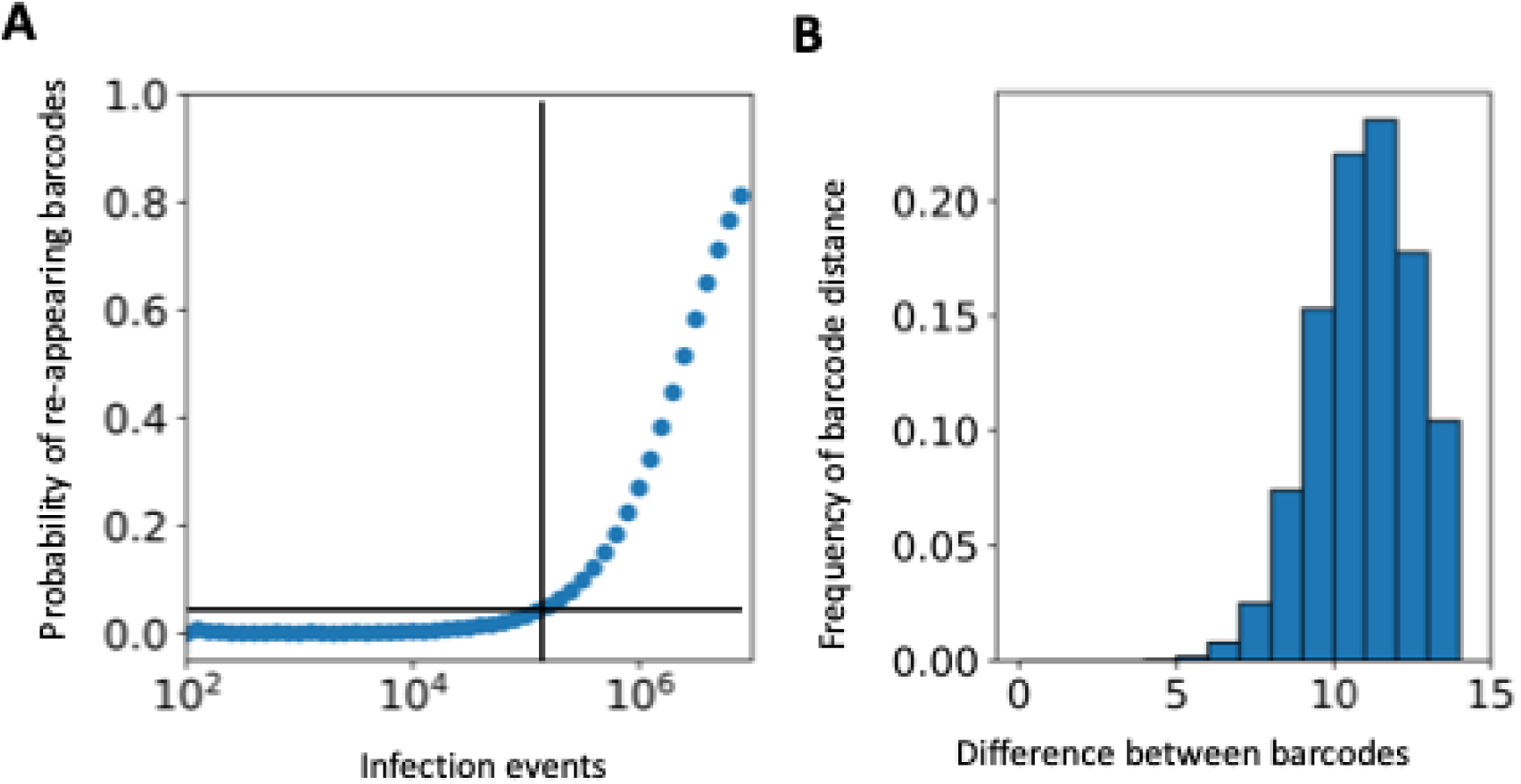
The quality controls of the barcode library. **A)** The probability of two cells sharing the same barcode with different infection size. Each dot is a simulation. X-axis is the number of infected cells. The y-axis is the probability of two cells are infected by the viruses having the same barcode. **B)** The histogram of the hamming distance (the number of different nucleotide) between any two barcodes.

**Figure S2.**
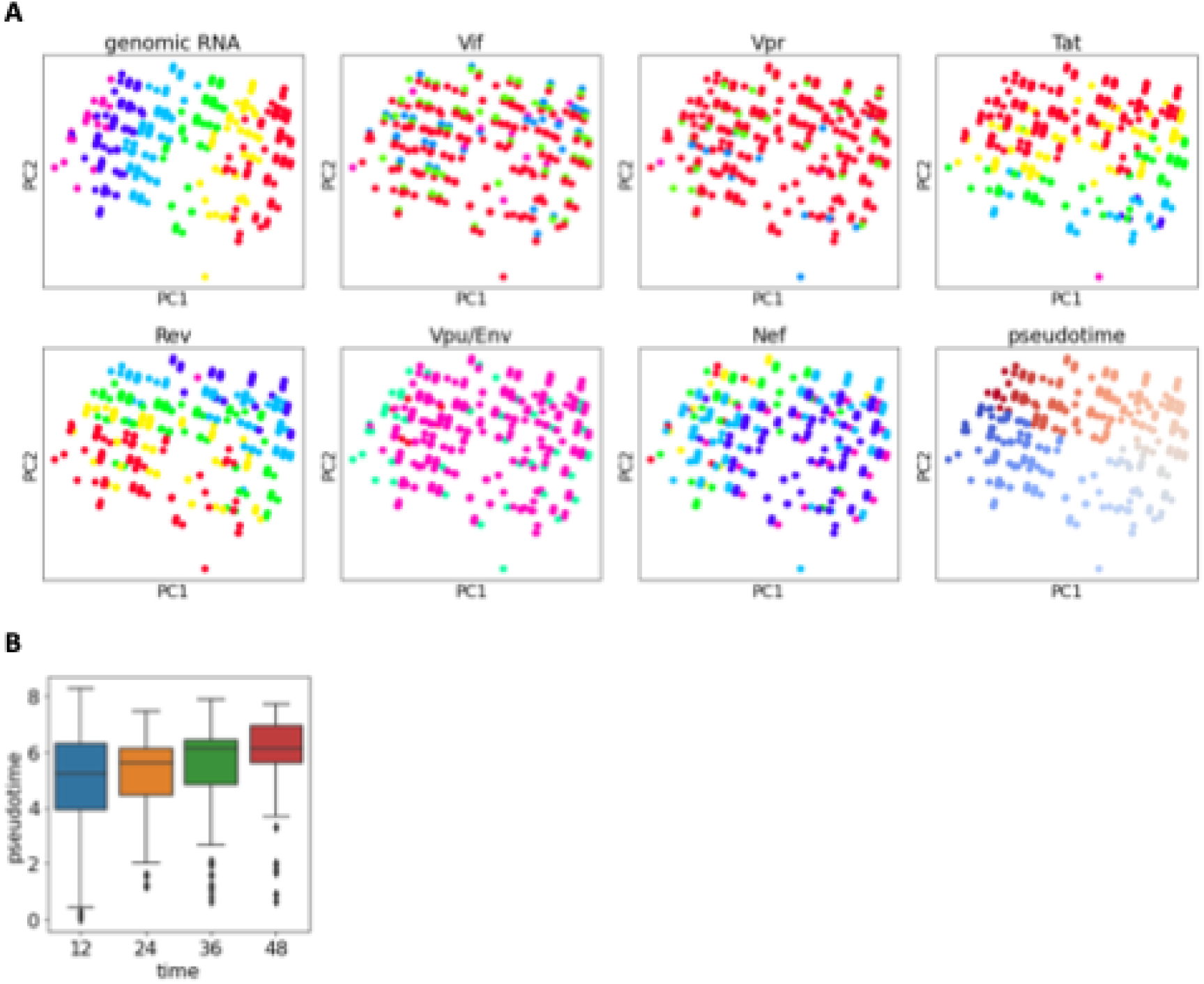
The PCA visualization of virus gene expression. **A)** The PCA visualization of virus gene expression, colored by gene abundance. **B)** The pseudotime inference of each barcoded virus.

**Figure S3.**
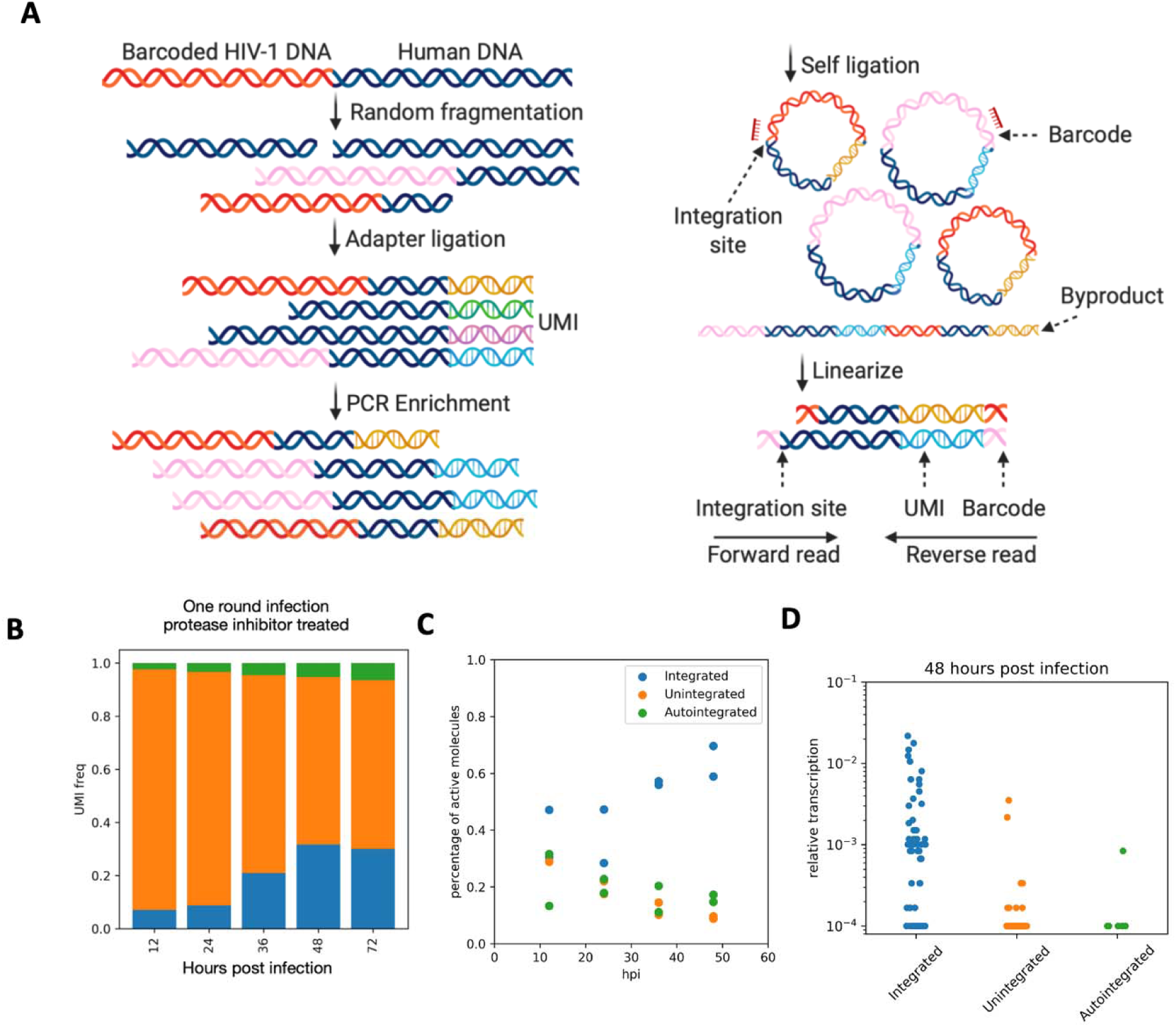
Transcriptional activity of different proviral conformations. **A)** Barcode - integration site linkage sequencing workflow. Genomic DNA was isolated from the infected cells. An L-shaped adapter was ligated to DNA fragments generated by enzymatic fragmentation. A 21-nucleotide UMI was included in the adapter to label each proviral molecule. Multiple steps of semi-nested PCR enriched the fragments containing the integration junction. The PCR product was digested and self-circularized to bring the barcode and the integration site into tandem proximity. Then the short fragments containing the UMI, barcode and the integration site were amplified for high-throughput sequencing. **B)** The abundance of different proviral DNA conformation. Primary CD4+ T cells were infected by the barcoded HIV-1 library. The total cellular DNA was extracted 12-72 hours post infection, and subject to barcode integration site linkage sequencing. The abundance of different proviral DNA conformation was quantified by analyzing the sequence at the integration junction. **C)** Percentage of active provirus at different time points. The proviruses with barcodes also observed in cellular RNA was defined as active provirus. Each dot represents one biological repeats. **D)** The relative transcription activity of different proviral conformations. The proviral transcription activity was calculated using the relative frequency of the barcodes in the cellular RNA. Each dot represents a barcode.

**Figure S4.**
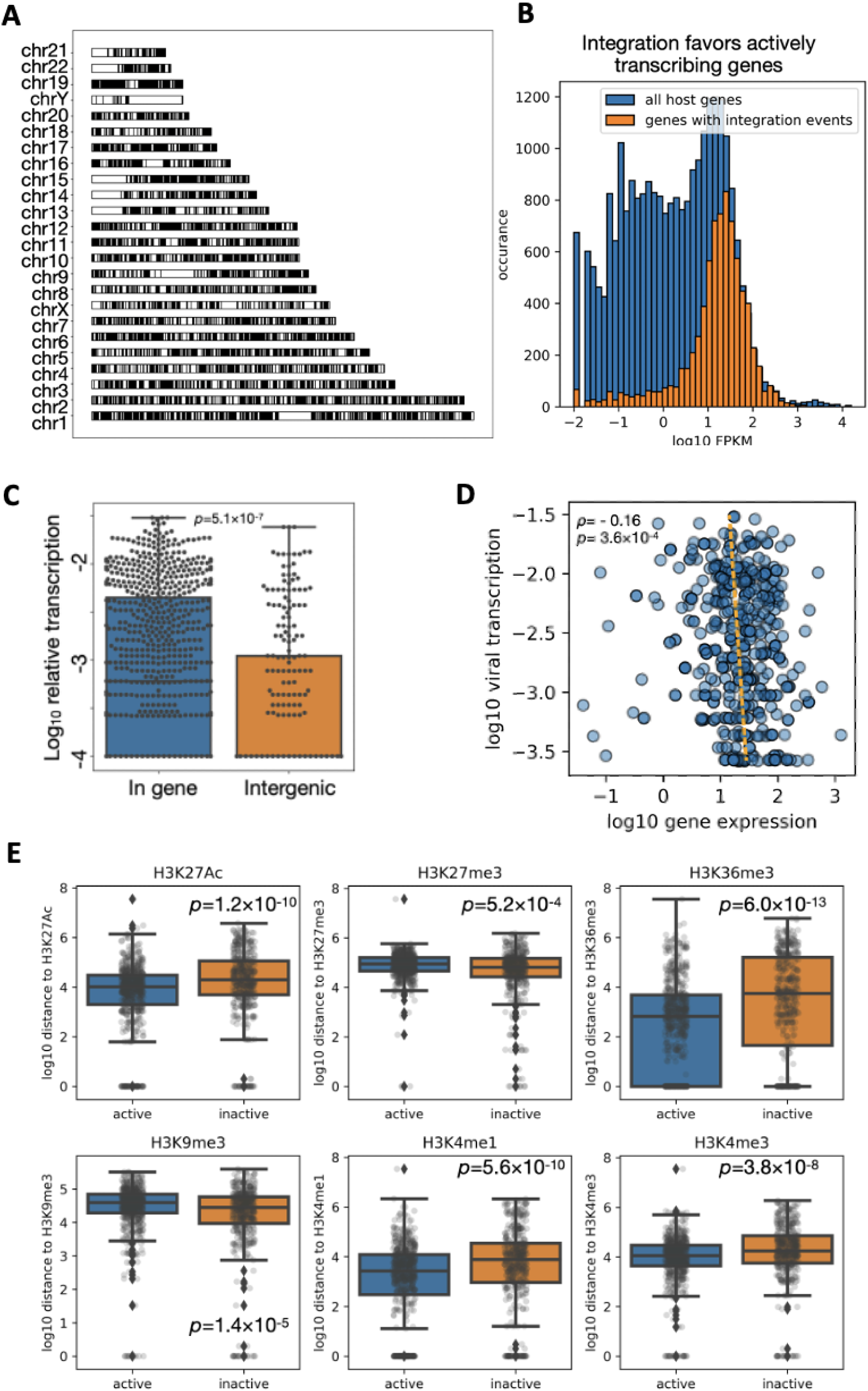
The positional effect of the HIV-1 integration site. **A)** The distribution of HIV-1 integration sites in primary CD4+ T cells’ chromosomes. The black bars represent the regions with integration sites observed in this study. **B)** The histogram of the host gene transcriptional activity. The host genes were classified according to whether there are proviral integration sites within their regions. The integration sites from Figure S3 are included for the analysis. **C)** The transcriptional activity of proviruses within or outside the gene regions. The transcriptional activity was calculated using the relative frequency of viral barcodes in cellular RNA. Wilcox’s rank sum test was performed. **D)** The correlation between the provirus transcription activity and the integrated host gene transcription activity. Each dot represents a provirus. The x-axis is the transcriptional activity of the host gene it integrated in. The y-axis is the transcriptional activity of the provirus. Spearman’s correlation test was performed. **E)** The distance to nearby histone modifications for active or inactive provirus. The provirus was classified as active and inactive according to whether its barcode was observed in cellular RNA. The distance between the integration site and the closet histone modification was shown. Each dot represents a provirus. ENCODE ChIP-seq data of human primary CD4+ T cells with H3K27ac, H3K27me3, H3K36me3, H3K9me3, H3K4me1 and H3K4me3 were used for the analysis. Wilcox’s rank sum test was performed.

**Figure S5.**
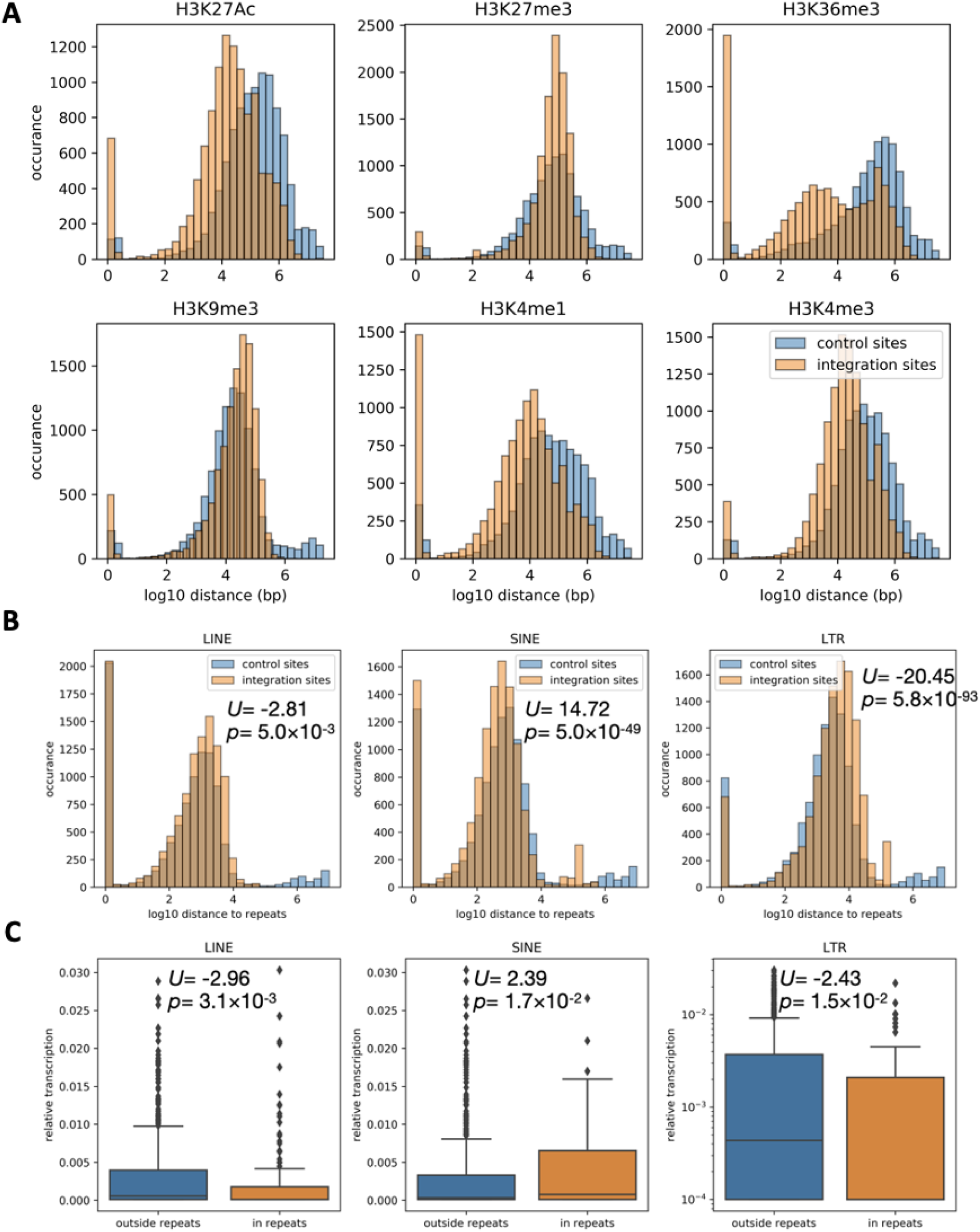
Distribution of integration site. **A)** The histogram of the distance between the integration sites to the nearest histone markers. **B)** The histogram of the distance between the integration sites and the nearest repeat regions. **C)** Transcriptional activity of provirus within repeat regions. The transcriptional activity of the provirus was quantified using the relative frequency of the viral barcode in the total RNA. The position of each integration site was compared with the LINE, SINE and LTR repeats annotation on the human genome. Wilcox’s rank sum test was performed.

**Figure S6.**
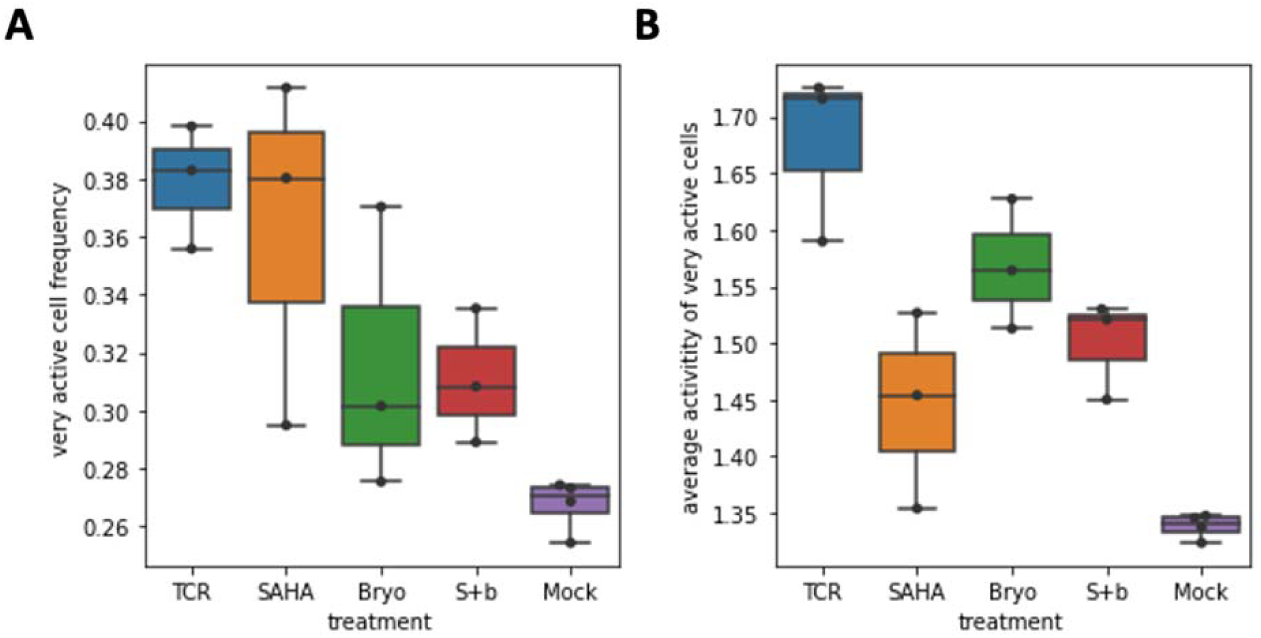
Transcriptional activity of LRAs treated cells. **A)** Frequency of active provirus. The provirus was classified as active and inactive according to whether its barcode was observed in total RNA. **B)** Transcriptional activity of the active provirus. Each dot represents an independent experiment.

